# Oxytocin reduces interoceptive influences on empathy-for-pain in the anterior insula

**DOI:** 10.1101/2021.10.22.465431

**Authors:** Sophie Betka, Cassandra Gould Van Praag, Charlotte L Rae, Gaby Pfeifer, Henrique Sequeira, Theodora Duka, Hugo Critchley

## Abstract

Empathy-for-pain states are underpinned by *interoception*, i.e the central representation of internal states. Cardiac signals occur in a phasic manner; baroreceptor discharges at systole communicate the heartbeats’ strength. These signals modulate pain and emotion processing. We tested whether these phasic interoceptive signals modulate empathy-for-pain. As oxytocin (OT) enhances empathy and modulates interoceptive signals’ precision, we also tested if OT administration impacts empathy-for-pain via interoceptive mechanisms.

Male subjects (N=32) attended three sessions to perform psychometric tests and an fMRI empathy-for-pain task, after intranasal administration of OT or placebo (40IU). Pictures of hands in painful or non-painful context were presented at systole or diastole. Effects of drug, emotion and cardiac timing on behaviour and brain activity was tested using general and mixed-effects linear models.

Across conditions, activation was observed within regions implicated in pain and empathy-for-pain, with insula activation greater in the right than left hemisphere. OT administration, compared to placebo, attenuated the reactivity of some regions, including anterior cingulate cortex, but presentation of stimuli at systole blocked the OT attenuating effect.

Our data suggest that OT alters the processing of motivationally-salient social cues, interacting with interoceptive signals. Our findings may inform targeted use of OT in psychiatric conditions linked to aberrant interoceptive processing.

## Introduction

Empathy (i.e. understanding the affective states of other people) is crucial to social cognition and is expressed for (higher-order) thoughts and emotional feeling states and for lower-level physical sensations, e.g. pain (Decety and Jackson, 2004, Singer *et al*., 2004, Kanske *et al*., 2017). Within the brain, the empathic processing of another’s pain engages a network of regions, encompassing dorsal anterior/middle cingulate cortex (ACC, MCC) and bilateral insulae (for review Lamm *et al*., 2011).Here, we refer to this network as the empathy-for-pain matrix.

Interoception, i.e. the sensing of visceral signals, has attracted a resurgence of interest, with interoceptive predictive coding is proposed as the mechanism underpinning emotions and self-representation (Seth, 2013). By extension, “interoceptive simulation” is hypothesised to underpin empathy: People with poor interoceptive skills (i.e. interoceptive accuracy, Garfinkel *et al*., 2015) have difficulty in identifying their own emotions, recognizing emotional facial expressions, or inferring the mental state of others (Terasawa *et al*., 2014, Betka *et al*., 2017, Bornemann and Singer, 2017, Shah *et al*., 2017). The amplitude of heartbeat-evoked potentials (a cortical signature of afferent interoceptive signals from the heart; Park *et al*., 2017) is associated with greater self-reported empathy (Fukushima *et al*., 2011). Moreover, people with better interoceptive accuracy show an enhanced capacity for imitating others, and an increased sensitivity / decreased tolerance to pain (Pollatos *et al*., 2012, Ainley *et al*., 2014).

These observations are consistent with the proposal that empathic understanding arises from a simulation (interoceptive prediction) of likely internal bodily state, requiring the integration of subsequent interoceptive afferent signals into emotional representations of both self and other. The insular cortex is proposed to be the neural substrate of such integration (Singer *et al*., 2009).

Interoceptive signalling, exemplified by the phasic firing of arterial baroreceptors with each heartbeat, influences our behavior (Łukowska *et al*., 2018). Within the brain, the magnitude of a preceding heartbeat-evoked potential, a putative cortical expression of baroreceptor activation, predicts the detection of a near-threshold visual stimulus (Park *et al*., 2014). In some instances, visual stimuli are harder to detect if they mirror the timing of heartbeats (Salomon *et al*., 2016). Insular activation mediates this effect, showing reduced activation to stimuli presented at the cardiac frequency. Indeed, there can be a cancelling out of external stimuli that appear ‘self-related,’ predicted by heartbeat frequency (Salomon *et al*., 2016). These self-related signals can enhance the attribution of self: For example, in the Rubber Hand Illusion, in which individuals are induced to perceive an artificial hand as their own, the effect of the experience is strengthened if the artificial hand pulses in synchrony with the participant’s heartbeat (Suzuki *et al*., 2013).

Cardiac interoceptive effects often relate to higher-order cortical processing, involving insula. However, a model dating back to the 1970s suggests that baroreceptor activation also has a lower-level inhibitory influence on sensory processing and cortical excitability (Lacey and Lacey, 1970, Walker and Sandman, 1979, Mini *et al*., 1995). The processing of pain stimuli is suppressed: Autonomic and insular responses to electrocutaneous shocks is attenuated if the shocks are delivered at cardiac systole (between 100 to 500ms after the electrocardiography (ECG) R wave, coincident with the peak of baroreceptor signaling(Eckberg and Sleight, 1992) (Gray *et al*., 2009, Gray *et al*., 2010).

In contrast, the perceived intensity of specific emotional facial expressions, of disgust and fear, can be enhanced when presented at systole, during natural baroreceptor activation (Gray *et al*., 2012, Garfinkel *et al*., 2014). Interestingly, more sustained artificial baroreceptor stimulation *via* neck suction, particularly on the right side, attenuates emotional ratings of fearful faces and inhibits activation within the insula, amygdala, hippocampus, thalamus, and brainstem (Makovac *et al*., 2015, Makovac *et al*., 2017). Taken together, these data link cerebral excitability to baroreceptor signalling.

Oxytocin (OT), a neuropeptide hormone, has a major role in mother-child bonding. OT also contributes more generally to prosocial behaviours across animal species (Kendrick *et al*., 1987, Carter, 2003, Young and Wang, 2004, Parker *et al*., 2005). In humans, intranasal administration of OT can increase interpersonal trust and increase attention toward social stimuli including faces (Guastella *et al*., 2008, Keri and Kiss, 2011). Moreover, OT increases sensitivity (recognition and detection) of facial emotions (Schulze *et al*., 2011, Leknes *et al*., 2013). Intranasal OT administration improves inferences about others’ social emotions, i.e. the cognitive component of empathy (Domes *et al*., 2007, Aoki *et al*., 2014) and, across individuals, the level of endogenous (salivary) OT correlates positively with both objective and subjective measures of emotional competence and interpersonal skills (Koven and Max, 2014). OT can also enhance aspects of empathy-for-pain (e.g. when adopting the other, but not self-, perspective) (Abu-Akel *et al*., 2015) and extend empathy beyond its usual in-group bias (Shamay-Tsoory *et al*., 2013). Two neuroimaging studies explore the impact of intranasal OT on vicarious pain processing: One study did not observe an OT-induced effect on neural correlates of empathy-for-pain, whereas the second study observed deactivation within secondary somatosensory cortices, insula, and MCC (Singer *et al*., 2008, Bos *et al*., 2015). One explanation for this discrepancy is that OT has intrinsic analgesic properties (Rash *et al*., 2014, Goodin *et al*., 2015, Tracy *et al*., 2015). Correspondingly, OT administration reduces subjective pain ratings and decreases the amplitude of pain-evoked potentials (Paloyelis *et al*., 2016). Moreover, intranasal OT enhances the inhibition of endogenous pain (Goodin *et al*., 2014). A second explanation is that OT modulates the impact of interoceptive signals, i.e. affecting the ‘visceral simulation’ component of empathy-for-pain. To date, there is limited empirical detail regarding the impact of OT on interoception. In rodents, OT injection has a direct impact on the nucleus of the solitary tract (NTS) (Karelina and Norman, 2009), facilitating visceral afferent synaptic transmission within the brain (Peters *et al*., 2008). Conversely, intranasal OT is observed to reduce objective measures of interoception on a heartbeat tracking task (requiring attention toward internal signals; Betka *et al*., 2018). Similarly, the OT-induced enhancement of right anterior insula activity has been found to correlate negatively with interoceptive accuracy scores (Yao *et al*., 2017). These observations are consistent with a model arising from a predictive coding framework, in which OT is proposed to modulate the precision of visceral representations (i.e. narrowing the attribution of salience to internal bodily signals). This OT neuromodulation biases attentional deployment toward relevant external cues (Ellenbogen *et al*., 2012), which in turn favours associative learning between internal and external cues, arguably a process fundamental to social cognition (Quattrocki and Friston, 2014).

In this study, we sought to characterise the impact of phasic visceral feedback on empathy-for-pain processing. Within the empathy-for-pain matrix, we expected attenuation at ventricular systole (during baroreceptor activity) relative to diastole (baroreceptor inactivity) when viewing of painful pictures compared to non-painful pictures.

The investigation of how interoceptive signals modulate the neural correlates of empathy further provides a novel opportunity to investigate the impact of intranasal OT on such processes. Specifically, we predicted that OT administration will modulate the impact of cardiac timing on empathy for pain, compared to placebo.

Based on earlier literature, we hypothesized that OT will decrease the activation of the empathy-for-pain matrix. However, taking into account the prediction that OT lowers interoceptive precision, we hypothesised that intranasal OT will reduce baroreceptor-related inhibition of cortical reactivity at systole. Therefore, activity within the empathy-for-pain matrix will be reduced by OT, but such effects will be primarily visible at diastole and attenuated by the systolic effects of baroceptor activation.

## Methods

### Participants

Thirty-two Caucasian male volunteers (mean age 25.1yrs; range 18–36yrs) took part in the experiment (see supplementary section for details regarding power calculation). Participants were recruited via advertisements at the University of Sussex and Brighton and Sussex Medical School. All participants were healthy individuals with no history of psychiatric or neurological conditions and were not taking medication. The average number of years of education was M=16.9 (SD=2.62). All participants gave written informed consent and were compensated for their time. The study was approved by the BSMS Research Governance and Ethics Committee.

### Procedure

The study was composed of three sessions. Participants were asked to abstain from drinking 24hrs before each session and were breathalysed for alcohol to confirm abstinence before the sessions. A urinary sample was collected to confirm the absence of drug use to exclude drug use disorder. Test results for all participants were confirmed as negative. During the baseline session, demographic data (e.g. age, education level) were recorded, a blood sample was collected and psychometric questionnaires were administered. A double-blind randomized within-subject design was used; the pharmacological sequence(OT/placebo) was randomly allocated to each participant for the second and third session. During these sessions, we computed the pulse transit time(PTT) for each participant, who then self-administered 40IU of OT nasal spray (Syntocinon;Novartis, Basel, Switzerland) or placebo (same composition as Syntocinon except for OT) in the presence of the experimenter (see supplementary section for details regarding the nasal spray administration). The participant transferred to the MRI scanner 5 minutes after administration. The second and third sessions were separated by 2-3 days. We failed to collect blood from two participants: to avoid omission of these participants in the behavioural analyses, we imputed the plasma OT missing data (see Supplementary Material). Plasma OT measures are detailed in Supplementary Material.

### Questionnaires

#### Toronto Alexithymia Scale-20 items (TAS-20)

The TAS-20 (Bagby *et al*., 1994) consists of 20 items rated on a five-point Likert scale (from 1 “strongly disagree” to 5 “strongly agree”), giving a total alexithymia score as the sum of responses across all 20 items.

#### Alcohol Use Questionnaire (Units of alcohol per week)

The AUQ (Mehrabian and Russell, 1978) is a 15-item scale measuring the quantity of alcohol consumption (alcohol units of 8g). Participants were asked to estimate the number of drinking days and the usual quantity consumed over the preceding six months. Here, we examined only the drinking quantity (i.e. alcohol units per week).

#### Trait Anxiety (STAI)

Trait anxiety was assessed using the Trait version of the Spielberger State/Trait Anxiety Inventory (STAI; Spielberger *et al*., 1983). This questionnaire is composed of 20 questions, such as “I lack self-confidence”. Participants answered each statement using a response scale (running from 1 “Almost never” to 4 “Almost always”) capturing a stable dispositional tendency (trait) for anxiety.

#### Beck Depression Index II (BDI)

Symptoms and severity of depression were evaluated using the BDI (Beck *et al*., 1996). Participants responded to 21 questions assessing the individual’s level of depression. The BDI items are scored on a scale from 0–3. All items were then summed for a BDI total score.

### Stimuli and Design

The empathy task consisted of sixty-four pictures illustrating a -Caucasian-hand, either in a painful or in a non-painful context (Jackson *et al*., 2005; see Figure 1.C). Each picture was presented twice, once at diastole and once at systole. Correspondingly, the design was a 2×2×2 repeated measures design with 2 levels of emotion (pain, no pain), 2 levels of cardiac cycle (systole/diastole), and 2 levels of drug (oxytocin/placebo). In total: 128 trials; 32 trials per condition; 4 conditions (pain /systole; pain/diastole; no pain/systole; no pain /diastole) by session. Importantly, participants were not aware of the cardiac timing manipulations.

**Figure 1.**
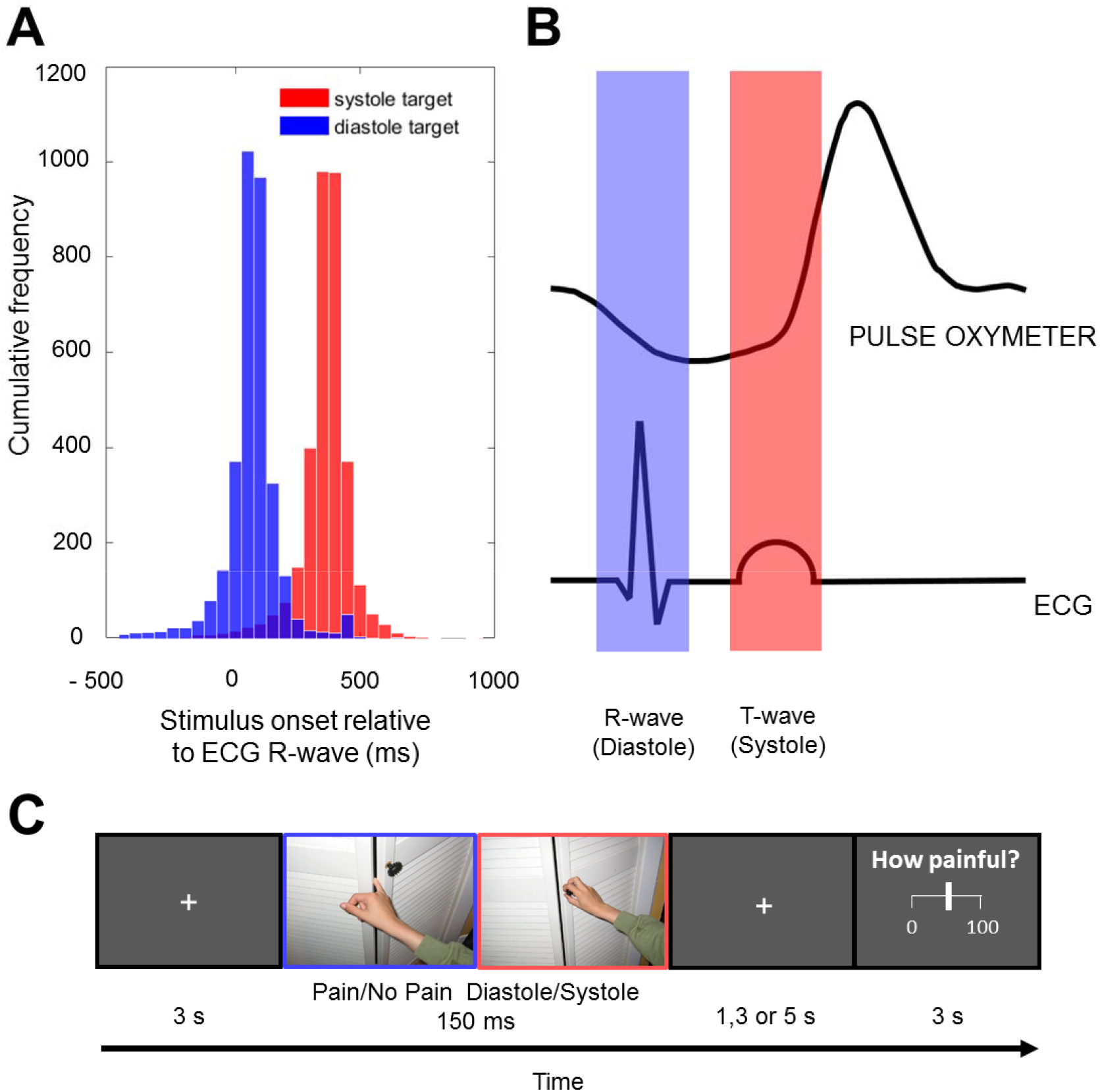
(A) Histogram detailing pictures presentation timing in relation to cardiac cycle within the MRI scanner. (B) Stimulus presentation was time-locked to coincide with two distinct points of the cardiac cycle: cardiac diastole in blue (ECG R-wave) and cardiac systole in red (ECG R-wave + 300ms, equivalent to ECG T-wave). (C) For the empathy-for-pain task, pictures of hand (painful context or non-painful context) were time-locked to diastole or systole. Participants made subsequent trial-by-trial pain intensity rating using a VAS.

### Empathy-for-pain task and cardiac timing

The empathy-for-pain task comprised the presentation of (1) a fixation cross (3000ms), (2) a picture (150ms) presented at either systolic or diastolic phase of cardiac cycle, followed by a jitter of 1000, 3000 or 5000ms, and (3) a visual analogue scale (VAS; 3000ms). During the VAS presentation, participants were asked to rate “How painful was the picture?”, using a scale of 0 (not painful at all) to 100 (very painful). See Figure 1A-B for a histogram detailing the precision of stimulus presentation in relation to cardiac cycle in the experiment conducted within the MRI scanner and Figure 1C for a depiction of the paradigm.

During the experiment, real-time cardiovascular timing information was obtained via an MRI-compatible pulse oximeter (8600Fo; Nonin Medical Inc., MN, USA) with the sensor attached to the participant’s left index finger. The peak of the finger pulse oximetry waveform (POW) reflects the R-wave delayed by the pulse transit time(PTT). By considering three previous interpulse intervals (generally called interbeat interval; IBI), we predicted the occurrence of the next finger pulse (R-wave + PTT) with validated accuracy (Gray *et al*., 2010, Gray *et al*., 2012). We subtracted the PTT from the predicted time of the next peak of POW, to derive the R-wave timing (coincident with diastole, i.e. baroreceptor quiescence). Similarly, subtracting the PTT from the predicted time of the next peak of POW and adding 300 ms, predicted the timing of ventricular systole i.e. high baroreceptor activation).

### MRI acquisition

Imaging data were collected using a 1.5 Tesla MRI scanner (Siemens Magnetom Avanto) with a 32-channel head coil. Visual stimuli were presented on a within-bore rear projection screen, at a viewing distance of approximately 45 cm. Stimuli were delivered using Cogent2000 v1.32 in MATLAB R2012b (The MathWorks, Inc., Natick, MA).

A T2*-weighted multiband echo-planar imaging(EPI) sequence was used, with a slice acquisition acceleration factor of 2. Each volume consisted of 36 axial slices oriented 30 degrees relative to the AC-PC line, covering the whole brain. Slices were acquired in an ascending and interleaved order. The following functional imaging parameters were used:TR=1379ms, TE=42ms, flip angle 90°, matrix= 64×64, FoV=192×192mm, slice thickness=3.0mm with a 20 % gap, resulting in 3.0 mm isotropic voxels. The exact number of fMRI volumes acquired depended on participants’ heart rate (mean volumes acquired:950). A T1-MPRAGE structural was acquired for registration (TR=2730ms, TE=3.57ms, 1×1×1mm resolution).

### Statistical analyses

#### Behavioural data

Analysis of pain ratings used a linear mixed-effects model, since the outcome was continuous, conducted with the lme4 package (Bates *et al*., 2015) in R (version 3.4.2; RCoreTeam, 2013). P values were computed using lmerTest package (Kuznetsova *et al*., 2014).

Ratings were analysed with drug (2 levels: Placebo=0; OT=1), emotion (2 levels: No Pain=0; Pain=1), Cardiac Timing (2 levels: Diastole=0; Systole=1) and their interactions, as fixed factors. The intercept reflected the average pain rating in the placebo no pain diastole condition. Participants were treated within the analysis as a random factor (see supplement for details).

This basic model was then compared to a similar model that also included anxiety, depression, alexithymia, alcohol intake, sequence, interbeat interval (IBI), fMRI jitter and plasma OT levels to test confounding effects of these variables. We contrasted the goodness of fit of these models using likelihood ratio tests.

To test for possible IBI changes due to OT administration, a mixed-effect linear model was fitted to the data, with IBI as outcome and drug as predictor. Participants were treated as a random factor, with random correlated intercepts and slopes, in function of the drug.

#### Neuroimaging data

The first ten images were discarded to allow the steady-state magnetisation. Images were converted from dicom to nifti format and re-aligned to anterior commissure. Images were slice-time corrected, spatially realigned to the first image, and unwarped using the acquired field and magnitude maps in SPM12. The T1 image was co-registered to the mean functional image and subsequently segmented to obtain normalisation parameters based on the standard MNI template. The segmentation parameters were used to transform each subject’s functional images and the bias-corrected structural image into MNI152 space. Voxel sizes of the functional and structural images were retained during normalisation, and the normalised functional images were spatially smoothed using an 8mm Gaussian kernel (full-width-half-maximum).

Task events were analysed in a general linear model, composed of two sessions (OT and Placebo). Each session included four regressors representing the onset and duration of the presentation of (1) painful images at systole, (2) painful images at diastole, (3) non-painful images at systole, and (4) non-painful images at diastole respectively. In addition, for each session, a further regressor, comprising the onsets and durations of VAS presentation, was added to the general linear model to separate the BOLD signal relating to pain rating from stimulus presentation. Movement regressors were included as confounds (6 head movement parameters calculated from scan volume realignment). Single-regressor T-contrasts were generated for each condition ((1) Oxytocin Pain Systole; (2) Oxytocin Pain Diastole; (3) Oxytocin No Pain Systole; (4) Oxytocin No Pain Diastole; (5) Placebo Pain Systole; (6) Placebo Pain Diastole; (7) Placebo No Pain Systole; (8) Placebo No Pain Diastole). These were entered into a full factorial second-level analysis, with drug, emotion and cardiac timing as non-independent (repeated measures) factors. Given that control variables did not improve the model’s fit at the behavioural level, we did not model them within the final full factorial design.

Contrasts were generated to test for (1) all experimental effects (F contrast: [8 experimental conditions]), (2) drug effects for both emotion and cardiac timing (Oxytocin Pain Systole; Oxytocin Pain Systole; Oxytocin No Pain Systole; Oxytocin No Pain Systole; Placebo Pain Systole; Placebo Pain Systole; Placebo No Pain Systole; Placebo No Pain Systole), (3) drug effect (Placebo>Oxytocin), (4) emotion effect (Pain>No Pain), and (5) cardiac effect (Diastole>Systole; T contrasts). In addition, the following contrasts were tested for interactions (PL(PvsNP)>OT(PvsNP); PL(PDvsNPD)>PL(PSvsNPS); OT(PSvsNPS)>OT(PDvsNPD)). Statistic images were thresholded at an initial cluster-forming threshold of p<0.001 for cluster-wise False Discovery Rate (FDR) correction for multiple comparisons at p<0.05 (Chumbley *et al*., 2010). Significant clusters were localised according to the Anatomy toolbox (v 2.2b, Eickhoff *et al*., 2007; see supplementary Table S1). Contrast estimate effect size plots were generated in SPM12 for each condition, at the peak coordinate of significant (empathy-for-pain related) regions according to the F test for all effects (Figure 3 & Figure 5).

## Results

### Psychometric, plasma and physiological measures

Means, standard deviations, and ranges of psychometric measures, basal plasma OT level as well as mean IBI were computed for sessions (Table 1).

**Table 1:**
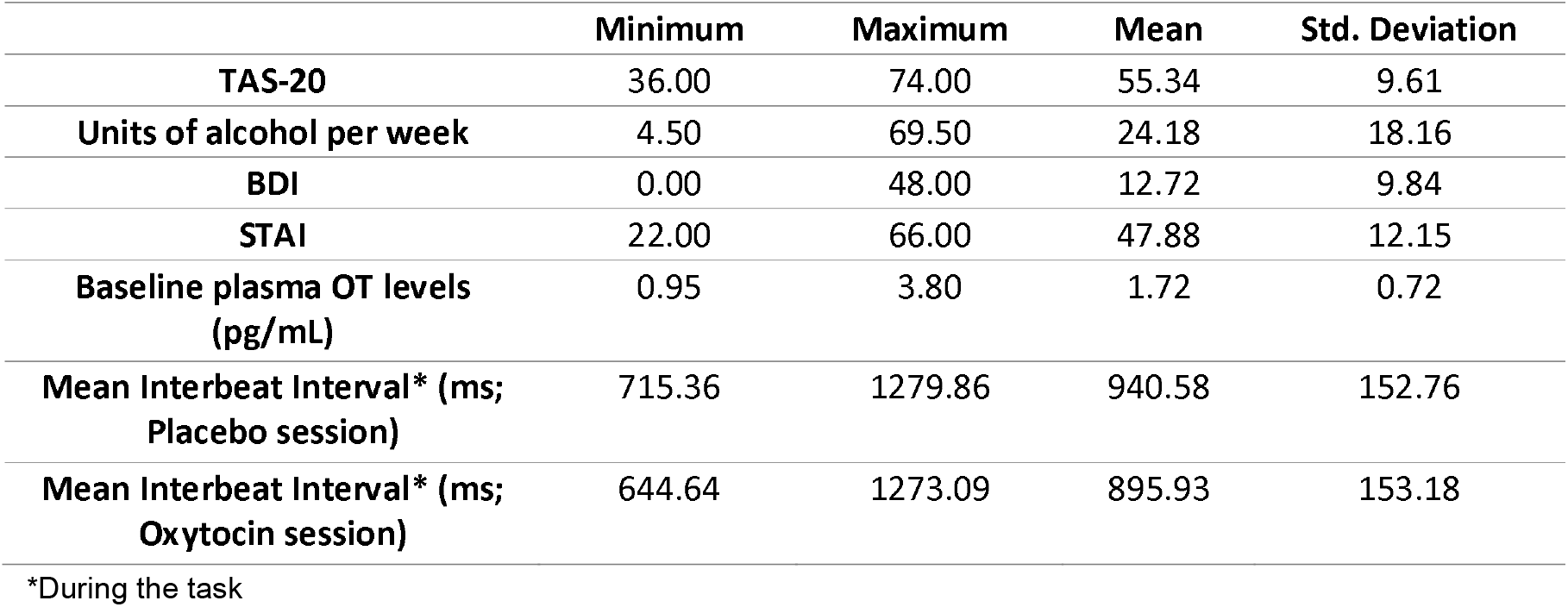
Psychometric measures

### Drug impact on cardiovascular activity during the task

A main effect of drug was observed on the IBI: OT reduced the IBI compared to placebo (i.e. increased heartrate; β=-44.66; SE=19.04; t=-2.346, *p*=0.0256, Table 2).

**Table 2.** Mixed-effects linear model assessing predicting the effect of oxytocin on interbeat interval (IBI) duration. The model includes IBI as outcome, drug as fixed factors, and participants as random factor, with random correlated intercepts and slopes, in function of the drug. The intercept reflects the average IBI in the placebo condition.

| | $\beta$ | SE | df | t | p | |
| --- | --- | --- | --- | --- | --- | --- |
| **Intercept** | 940.589 | 27.005 | 31.003 | 34.831 | 0 | *** |
| **Oxytocin** | -44.656 | 19.037 | 31.013 | -2.346 | 0.026 | * |

### Behavioural performance

A main effect of emotion was significant. Painful pictures were rated as more painful than non-painful stimuli (see **Error! Reference source not found**.; *p*<0.001, Table 3). No suprathreshold effect of drug or cardiac timing was observed. The addition of control variables (anxiety, depression, alcohol units per week, alexithymia, sequence, IBI, endogenous plasmatic OT or jitter) to the model did not significantly improve the goodness of fit (Basic model: AIC=72 103; Model with covariates: AIC=72 106; comparison: χ^2^(8)=13.402; *p*=0.098).

**Table 3.** Mixed-effects linear model predicting changes in pain rating. The model includes drug (2 levels: Placebo=0; OT=1), emotion (2 levels: No Pain =0; Pain=1), Cardiac Timing (2 levels: Diastole=0; Systole = 1) and their interactions, as fixed factors; participants, as random factor The intercept reflects the average pain rating in the placebo no pain diastole condition.

| | $\beta$ | SE | df | t | p | |
| --- | --- | --- | --- | --- | --- | --- |
| **Intercept** | 19.363 | 2.475 | 34.392 | 7.823 | 0.000 | *** |
| **Drug** | -0.267 | 1.371 | 67.468 | -0.195 | 0.846 |  |
| **Emotion** | 43.582 | 0.905 | 8046.889 | 48.169 | 0.000 | *** |
| **Cardiac** | 0.89 | 0.903 | 8047.142 | 0.985 | 0.324 |  |
| **Emotion\*Cardiac** | 1.661 | 1.273 | 8046.869 | 1.305 | 0.192 |  |
| **Drug\*Cardiac** | -0.443 | 1.272 | 8046.997 | -0.348 | 0.728 |  |
| **Emotion\*Cardiac** | -1.853 | 1.278 | 8047.023 | -1.449 | 0.147 |  |
| **Drug\*Emotion\* Cardiac** | 0.629 | 1.8 | 8046.936 | 0.35 | 0.727 |  |

### Neuroimaging

Activity associated with viewing of pain and no pain stimuli, presented at systole and diastole in both drug conditions, was examined using an F contrast testing for all experimental effects. Activations were observed across a canonical set of pain processing regions, including inferior, superior and middle frontal gyri, precentral and postcentral gyri, superior parietal lobule, cerebellum and precuneus. This network encompassed activation of the empathy-for-pain matrix, engaging anterior insulae (AI; extending to inferior frontal gyri); anterior and middle cingulate cortex (ACC/MCC). Also, this network encompassed implicated in the representation of information concerning body parts, e.g. fusiform gyrus, (Figure 3A, Supplementary Table S1).

All whole-brain contrasts testing for drug, emotion, cardiac differences or interactions did not attain significance at corrected thresholds.

Examining the experimental effects for emotion and cardiac timing for OT and placebo (Oxytocin Pain Systole; Oxytocin Pain Diastole; Oxytocin No Pain Systole; Oxytocin No Pain Diastole; Placebo Pain Systole ;Placebo Pain Diastole; Placebo No Pain Systole; Placebo No Pain Diastole; Figure 2B, D, F, H, C, E, G, and I respectively; Supplementary Table S1) revealed activation in both drug conditions across precuneus, superior parietal lobule, superior frontal gyrus, lingual gyrus, occipital and frontal poles, cerebellum, as well as putamen. While bilateral AI were activated for all experimental conditions, AI showed greatest activation during the pain systole condition. Activation of the ACC was generally attenuated by OT, except during the pain systole condition. This effect was not observed under placebo. Direct testing revealed that ACC was deactivated by OT, relative to placebo. Similarly, OT also deactivated paracingulate cortex, supplementary motor area, superior frontal gyrus and lateral occipital cortex (Figure 4; Supplementary Table S1.J).

**Figure 2.**
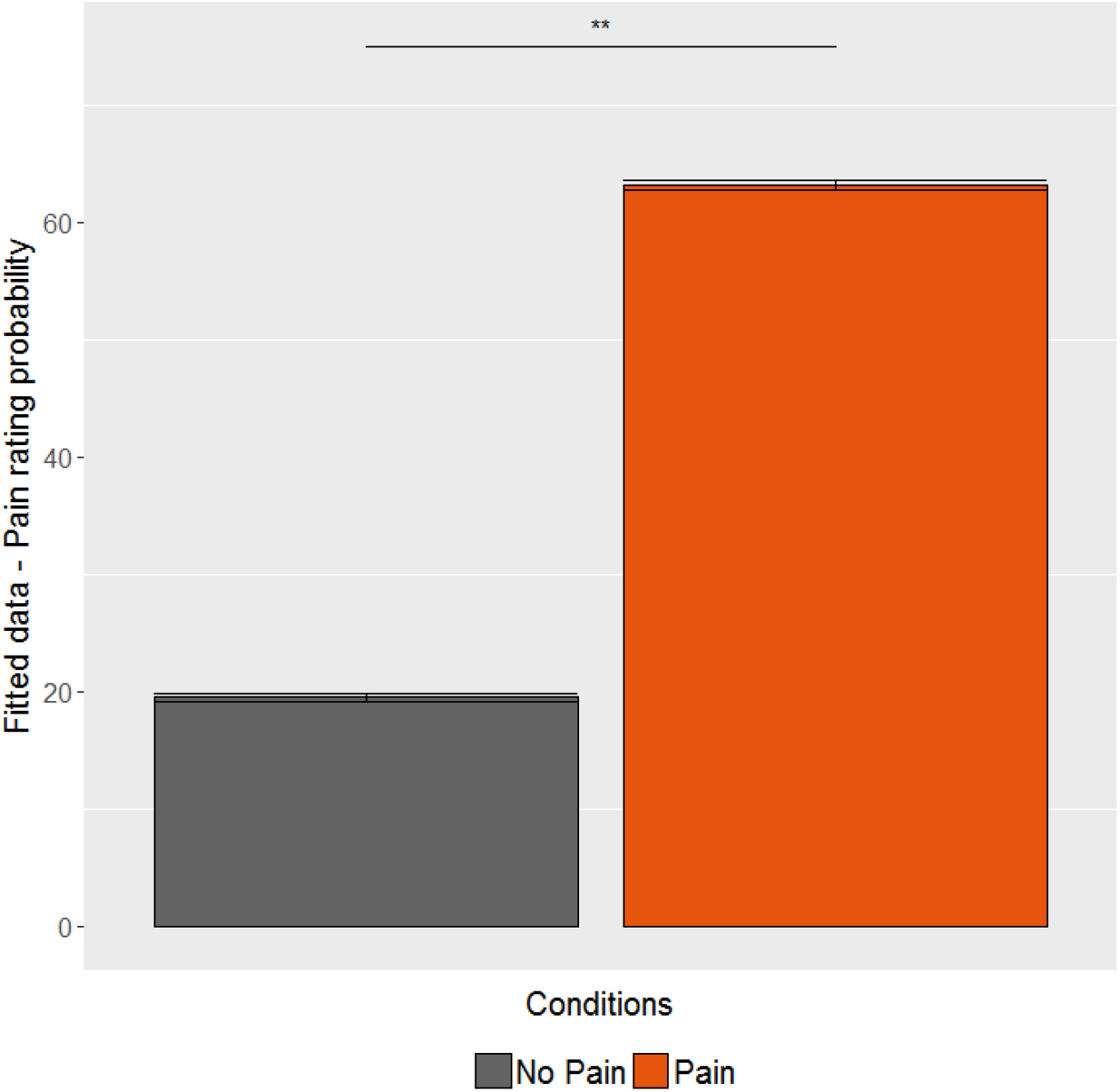
Bar plot with error bars (confidence interval) demonstrating the pain effect of emotion on pain rating: pain stimuli are rated more highly than no pain (p < 0.01).

**Figure 3.**
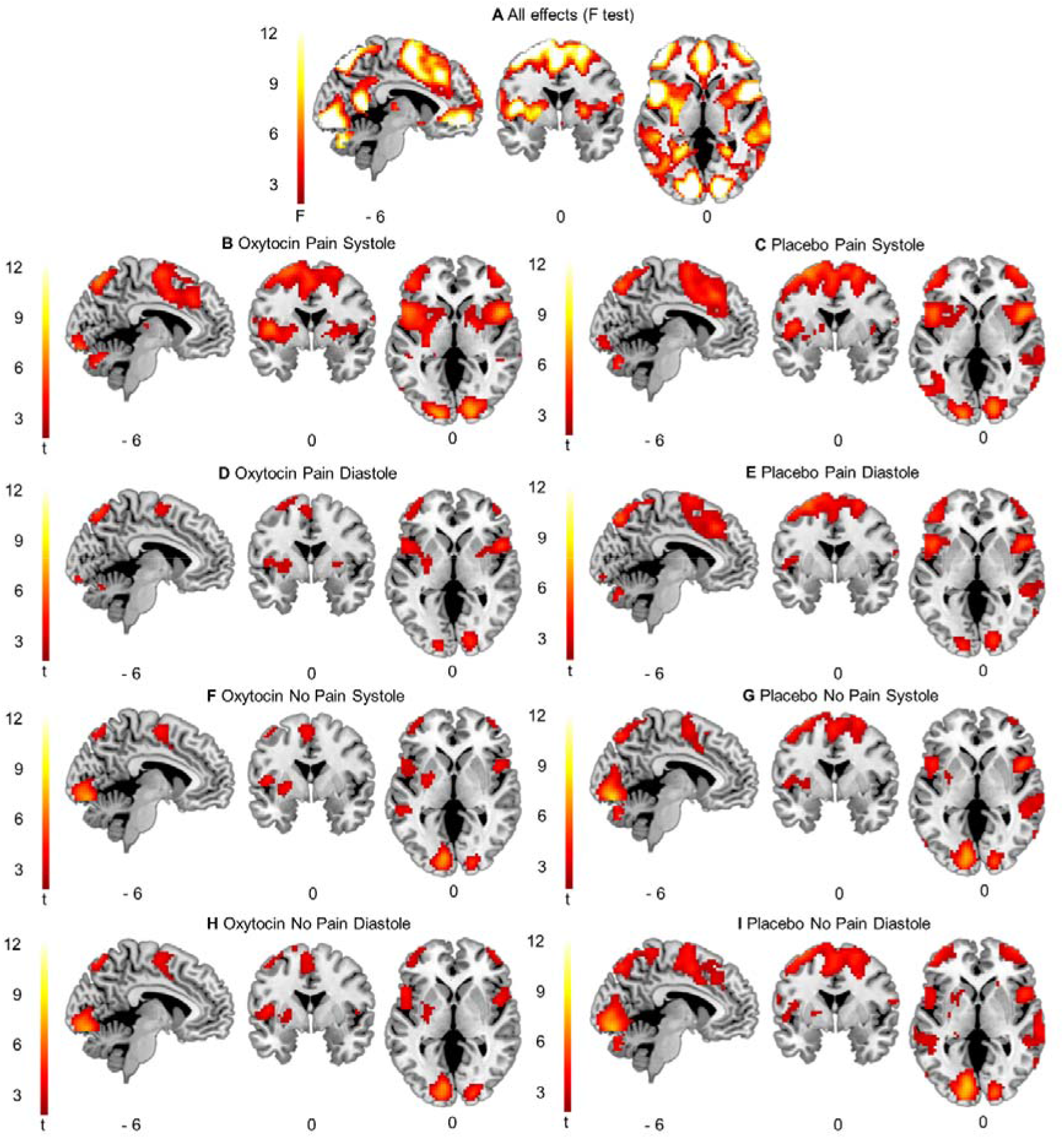
Activity during viewing pictures across experimental conditions. Oxytocin globally reduces the cerebral activation compared to placebo, except in the pain systole condition. (A) F test of all experimental effects, (B) Oxytocin Pain Systole; (C) Placebo Pain Systole; (D) Oxytocin Pain Diastole; (E) Placebo Pain Diastole; (F) Oxytocin No Pain Systole; (G) Placebo No Pain Systole; (H) Oxytocin No Pain Diastole; (I) Placebo No Pain Diastole. All images thresholded at cluster-wise FDR p < 0.05 (cluster-forming threshold p < 0.001).

**Figure 4.**
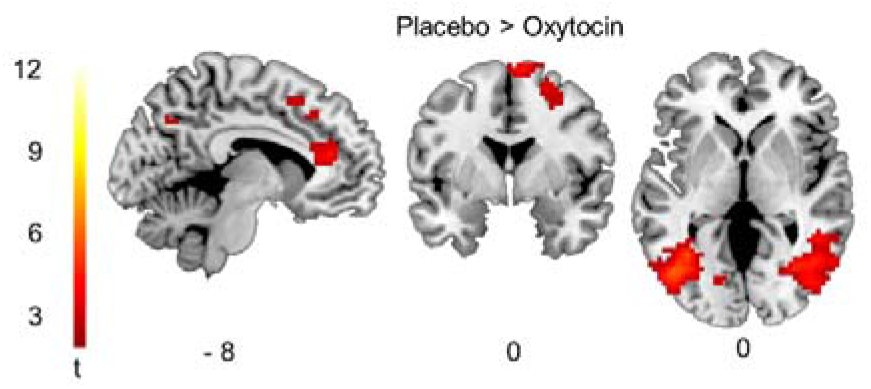
Activity during all experimental conditions under placebo minus all experimental conditions under oxytocin thresholded at cluster-wise FDR p<0.05 (cluster-forming threshold p<0.001).

Contrast estimate effect size plots, representing activity at the peak coordinate of bilateral AI and bilateral ACC clusters, identified according to the F test for all experimental effects, are presented in Figure 5. These reveal generally greater activity within right AI compared to the left AI, across all experimental conditions (Figure 5A&B). Under placebo, left AI showed greater activation for pain versus no pain condition, irrespectively of the cardiac timing. However, within right AI, this differential response between pain and no pain was less evident. This suggests that in the right AI, pain induced more activation than non-pain, and systole induced more activation than diastole. Strikingly, under OT, activation in bilateral AI was reduced relative to placebo in all conditions, except during pain systole, where the activation seemed preserved or even increased. In line with the right AI, under OT, activation in bilateral ACC was reduced relative to placebo (Figure 5C&D), except in the left ACC during pain systole, where the activation is preserved, or even increased (Figure 5C). However, these trends did not survive stringent significance threshold testing as formal interactions in whole-brain contrasts.

**Figure 5.**
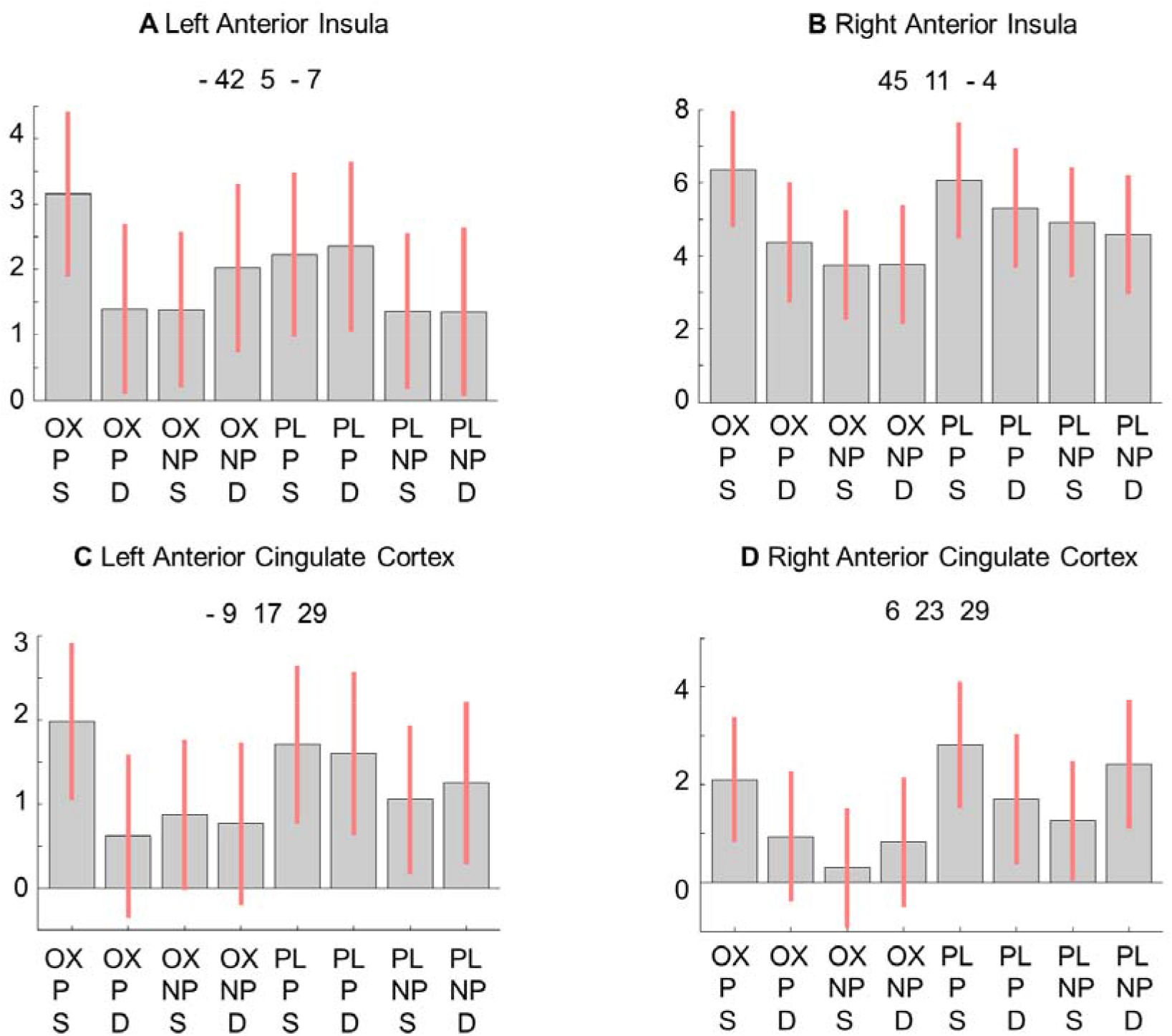
Contrast estimate effect size plots for the bilateral Anterior Insulae (A-B), and bilateral Anterior Cingulate Cortices (C-D), respectively, for (left-to-right) Oxytocin Pain Systole (OXPS); Oxytocin Pain Diastole (OXPD); Oxytocin No Pain Systole (OXNPS); Oxytocin No Pain Diastole (OXNPD); Placebo Pain Systole (PLPS); Placebo Pain Diastole (PLPD); Placebo No Pain Systole (PLNPS), and Placebo No Pain Diastole(PLNPD).

## Discussion

In the present study, we first sought to characterise the impact of (phasic) interoceptive cardiac signals on empathy-for-pain processing. In addition, we investigated a novel question related to the impact of intranasal OT on the relationship between visceral feedback and empathic brain activations. We found that momentary states of cardiovascular arousal, occurring at systole when baroreceptors are active, are associated with decreases in activity when participants have received intranasal oxytocin within two key pain-matrix regions: the right anterior insula, and anterior cingulate cortex. Although we observed effects were predicted, they did not attain threshold significance after stringent corrected for multiple comparison in whole brain imaging analyses. Nevertheless, three important deductions can be made. The right anterior insula (AI) appears to be specifically engaged in integrating emotional processing with interoceptive signals (associated with cardiac timing). Intranasal OT was observed to reduce brain activation generally across experimental conditions. However, it its notable the activation associated with processing painful pictures at cardiac systole remained unaffected by OT administration.

We observed a main effect of viewing the empathy-for-pain task stimuli, irrespective of cardiac timing or drug condition: Activity was elicited across cerebral areas previously implicated in the processing and perception of pain. This empathy-for-pain matrix encompassed bilateral AI and anterior/middle cingulate cortices (ACC/MCC). These areas also play a crucial role in salience attribution, a related factor accounting for activation of these brain areas during our task. Indeed, participants had to pay attention to pictures, which were briefly presented. As part of the “salience network”(Seeley *et al*., 2007), the AI is hypothesized to detect and prioritise the most relevant bottom-up events among internal and external environments, and to initiate and direct attentional control, which can be sustained by ACC (Menon and Uddin, 2010). Interestingly, activation of right AI was generally greater compared to the left AI, across all conditions. These data reinforce the salience hypothesis, as switching between central-executive and default-mode networks involves specifically the right AI (Sridharan *et al*., 2008).

In addition, the engagement of right AI in our experiment is likely to encompass the monitoring and appraisal of predicted visceral inputs, constituting a potential basis for selfhood (Craig, 2002, Critchley, 2004, Critchley *et al*., 2004, Critchley and Seth, 2012, Babo-Rebelo *et al*., 2016). Posterior insular cortices bilaterally receive an initial representation of interoceptive information or physical pain, it is proposed that the second-order interoceptive representation is constructed within right AI, before being projected on more frontal areas(Craig *et al*., 2000, Brooks *et al*., 2002, Craig, 2002). However, before allowing the necessary switching between main cerebral networks, internal physiological inputs received by the (right) AI need to be integrated with representations of external sensory information(Singer *et al*., 2009). Recently, a growing body of evidence supports the crucial role of right AI in these integrative processes(Garfinkel *et al*., 2014, Salomon *et al*., 2016, Salomon *et al*., 2018). Consequently, *via* our manipulation of stimulus presentation on and off the heartbeats, we engaged mechanisms enabling both interoceptive/exteroceptive integration and associated switching of cognitive resources, processes in which the right AI is specifically implicated.

Our second key finding was the OT-induced attenuation of cerebral activation: Across all experimental conditions, intranasal OT, relative to placebo, was associated with decreased activation over cerebral areas including paracingulate cortex, supplementary motor area, superior frontal gyrus and lateral occipital division. The reduction in the reactivity of the ACC by OT was particularly noteworthy. This result extends evidence showing that intranasal OT suppresses anterior cingulate activity when processing emotional faces (Labuschagne *et al*., 2012, Kanat *et al*., 2015, Luo *et al*., 2017) and reduces middle cingulate activation during empathy-for-pain (Bos *et al*., 2015). Both ACC and MCC (regions of dorsal ACC) are characterized by the presence of Von Economo neurons; large projection neurons from cingulate, AI and more frontal regions, observed in some mammals (Allman *et al*., 2011). Based in part on their concentrated presence in humans and great apes, it is postulated that these specialised pyramidal neurons play an important role in salience attribution, social cognition and even consciousness(Critchley and Seth, 2012, Evrard, 2019). The dorsal cingulate, including MCC, is connected to cognitive and motor-related areas, as well as thalamic nuclei and spinal cord. Rostrally, the ACC is increasingly also connected to cortical and subcortical structures involved in motivation behaviour and emotional responses(Stevens *et al*., 2011). However, both these regions react to unpleasant/noxious stimulation (Vogt, 2005, Shackman *et al*., 2011). Indeed, ACC encodes the motivational and affective dimension of pain, preparing behavioural responses to pain and other negatively-valenced stimuli (Morrison *et al*., 2004). Consequently, reaction times are speeded when faced with a potentially noxious challenge(Morrison *et al*., 2007). It is possible that the OT-induced decreased activation in the ACC might compromise the preparation and production of defensive actions. This hypothesis is consistent with observed analgesic effects of OT, modulating pain threshold and discomfort (Ohlsson *et al*., 2005, Rash *et al*., 2014), and corresponding autonomic reactions coordinated through ACC activation (Critchley *et al*., 2003). Therefore, the observed ACC deactivation might be a signature of reduced sympathetic activity after OT administration (Norman *et al*., 2011).

Our third finding is that while OT reduced activation within the empathy-for-pain matrix, but this effect was blocked when viewing pain stimuli at systole. At ventricular systole, arterial baroreceptors are activated by phasic ejection of blood from the heart (Bronk and Stella, 1932). A suppression of cortical excitability specific to baroreceptor activation is extensively described in the literature (Lacey and Lacey, 1970, Droste *et al*., 1994, Angrilli *et al*., 1997, Edwards *et al*., 2001). Indeed, presentation of stimuli time locked with baroreceptor activation decreases the activation of the insula, via the interoceptive pathway(Gray *et al*., 2009, Makovac *et al*., 2015). Stimuli occurring at the same frequency as the heartbeat are also inhibited (Salomon *et al*., 2016). Interestingly, there is evidence suggesting that OT elicits a reduction in interoceptive processing. Indeed, intranasal OT administration can decrease visceral symptoms in patients with irritable bowel syndrome (Louvel *et al*., 1996) and reduce objective performance accuracy on a heartbeat tracking task, via modulation of right AI activation (Yao *et al*., 2017, Betka *et al*., in press). These observations are corroborated by work in animals showing that OT can reduce sensitivity to bladder distension (Black *et al*., 2009, Engle *et al*., 2012). One model proposes that OT modulates interoceptive precision, facilitating attention deployment toward external cues and enhancing associative learning between internal and external stimuli (Quattrocki and Friston, 2014). By extension, OT through amplification of interoceptive signal-to-noise may enhance the processing of baroreceptor inputs, in the context of its usual suppression of cortical excitability. This effect would benefit the processing of emotional and salient external cues. Correspondingly, a clear pattern of preserved insular and cingulate activation during pain stimulation at systole was observed after OT administration. Further studies involving pharmacological manipulation and connectivity analyses might usefully explore this potential mechanism.

The present study should be considered in light of several constraints. First, our empathy-for-pain paradigm used pictures of hands in painful and non-painful contexts. A more efficient way to tap into empathic cerebral responses of vicarious pain could have used more explicit cues informing about another’s affective states. Indeed, this kind of paradigm elicits greater activation in cerebral structures associated with inferring and mentalizing other’s mental states (Lamm *et al*., 2011). A second limitation was that we did not screen our sample for atypical/ variant pain processing, notably mirror-sensory or mirror-pain synaesthesia(Grice-Jackson *et al*., 2017a). Moreover, different subpopulations of ‘pain responders’ show distinct patterns of functional neural connectivity between AI and the temporoparietal junction (Grice-Jackson *et al*., 2017a, Grice-Jackson *et al*., 2017b). Finally, there are methodological considerations: The time constraint inherent to the combination of pharmacological manipulation and neuroimaging, we did not measure change in PTT after the drug administration. It is possible that OT modulated PTT, which may have affected cardiac-contingent stimulus presentation. However, PTT correlates with systolic blood pressure, which is not modulated by acute intranasal OT(Rash and Campbell, 2014). Moreover, multiband scanning at a magnetic field of 1.5T constrained the statistical power of our study. Nevertheless, a clear pattern of preserved activation during pain stimulation under OT, at systole, emerged for AI and ACC. Higher MRI field strength may add further detail regarding wider neural correlates of this effect. Finally, one analytic model (factorial) was selected among other potentially appropriated approaches, as is often the case for sophisticated experimental designs.

Our findings support a mediating role of right AI in interoceptive monitoring and salience attribution. Even so OT seems to be a facilitator of empathy, yet, our data extend previous empirical evidence OT-induced neural deactivation and disengagement of the empathy-for-pain matrix. Finally, a clear pattern of preserved activation of the insular and cingulate cortex emerged under OT, during pain stimulation at systole. Taken together, our data suggest that OT alters the relationship between afferent interoceptive signals and the processing of relevant external cues. On a more hypothetical note, OT might play a key role in homeostasis mentalization as described by Fotopoulou and Tsakiris (Fotopoulou and Tsakiris, 2017). Psychiatric populations characterized by enhanced interoceptive signals could potentially beneficiate from therapeutic use of OT.

## Supporting information

Supplementary materiel

## Abbreviation

ACC: Anterior Cingulate Cortex;
AI: Anterior Insula;
MCC: Middle Cingulate Cortex;
POW: Pulse Oximetry Waveform;
PTT: Pulse Transit Time;

## Acknowledgements

We thank Prof. Sarah Garfinkel and Dr. Yannis Paloyelis for their assistance and advice.

