## Supplementary materiel for "Oxytocin reduces interoceptive influences on empathy-for-pain in the anterior insula"

### Supplementary material

### Power calculation

The sample size selected followed discussion with, and the advice of, a professional statistician. At the time of the creation of the project, no study looked into the impact of oxytocin on interoception. Therefore, we decided to use publications related to alexithymia, a condition related to a failure in interoceptive processing (Helling, 2009). A sample size of 30 was determined on the basis that it gave sufficient power to generate parameter estimates for the target measures that can be used alongside existing datasets. Moreover, equivalent group sizes were used in published studies from which we derived comparative data (Luminet *et al.*, 2011, de Haan *et al.*, 2014) facilitating comparison and inference across literature. We calculated that with n=30, assuming a correlation of 0.6 for within-subject measurements, the equivalent sample size for a parallel group trial would be 150 (i.e. 75 per condition). The minimum effect size we could detect with 80% power at p<0.05 significance would be 0.46. Assuming a standardized difference of 12.9, as per the De Haan’s paper, the minimum difference we could detect would be 6 points on the TAS-20 (alexithymia questionnaire). Therefore, with 30 patients in a within-subject design, we can detect a 6 point difference in TAS-20 with 80% power at p<0.05 significance assuming a standardized difference of 13 points and a within-subject correlation of 0.6. One of the main limitations of this approach was that we considered two behavioural studies in order to compute the sample size of a multimodal project (i.e. including neuroimaging, electrophysiology and behavioural measures). However, the expectation is that neuroimaging is more sensitive than behaviour, and thus, our power analysis valid for this current project.

### Experimental Procedure

In the present study, we were interested in the impact of intranasal oxytocin (OT) on interoception on empathy-for-pain. In order to take into account, the potential interindividual differences in endogenous OT levels of our participants, we measured plasma OT to include as a control variable in our analyses.

#### Plasma oxytocin (OT) measure

For the OT radioimmunological measurements, 10 ml blood was drawn into an ethylenediamine tetraacetic acid (EDTA) vial. Samples were mixed briefly and were then centrifuged in a refrigerated 4 °C centrifuge at 4° C for 10 min at 1300 g. A quantity of 0.8 ml plasma was pipetted into 2-ml Eppendorf vials and samples were kept at -80° C until shipping. OT specific radioimmunoassays were conducted by Professor Rainer Landgraf’s team at the University of Munich (http://www.riagnosis.com). Assays were strictly standardised and validated in animal and human studies across different physiological states (hypertonicity, parturition, lactation, stress, etc.) to reliably detect the bioavailable neuropeptide in peripheral (plasma) compartments (for more details on the procedure see Landgraf, 1985, Wotjak *et al.*, 1998, Kagerbauer *et al.*, 2013, Striepens *et al.*, 2013).

#### Nasal spray administration

1) Instructions related to the administration: The participant was advised that she/he will have to take 10 puffs of nasal spray: 5 in each nostril, right and left sides were alternated. As puffs needed to be spaced out, the experimenter gave a signal to the participant every 30 seconds, prior to each puff.

2) Instructions related to the nasal spray squeezing: Prior to the administration, participants were given a spray bottle full of water and were trained to deliver nice puffs in the air.

3) Instructions related to the nasal spray sniffing: The participant was instructed to give a quick and strong squeeze while quickly sniffing. The experimenter demonstrated the process as an example.

4) Instructions related to head position: Afterward, the experimenter demonstrated to the participant the head position she/he should take the nasal spray. “Put the nozzle in the nostril as far as you can. Do not touch the septum or the walls of your nose. Your head needs to be slightly back. The nozzle needs to be in line with your nose, no angle.”

When the participant felt ready, the administration began and lasted for 5 minutes in the presence of the experimenter. Nostril side sequence (left or right side first) was randomized across participants.

### Missing oxytocin plasma levels

Plasma OT levels were missing for two participants from whom we were unable to collect blood samples. Generalized and linear mixed models omit cases with missing data. Rather than losing two participants, we performed multiple imputation using predictive mean matching. Predictive mean matching (PMM) is a semi-parametric imputation approach which imputes missing values by means of the nearest-neighbour donor, with distance based on the expected values of the missing variables conditional on the observed covariates (Little, 1988). To do so, we used the Multiple Imputation by Chained Equations (MICE) package in R, with PMM as method of imputation and the number of imputations set at 5 (Van Buuren and Groothuis-Oudshoorn, 2011). Imputed data were checked and included in the dataset.

### Statistical Analyses

#### Random effect structure

We used a data-driven approach to specify the random effect structure (Barr *et al.*, 2013). Models were run that included (1) random intercepts and (2) random correlated intercepts and slopes. The goodness of fit of these models were compared using likelihood ratio tests. The model including random correlated intercepts and slopes explained more variance than the other models (Model 1: AIC =-72 202; Model 2: AIC =-72 103; Models 1-2 comparison: χ^2^ _(2)_ = 103.03; *p* < 0.001).

### Supplementary results

**Supplementary Table 1.** Local maxima of significant clusters per contrast, localised according to the Anatomy toolbox (V2.2b; Eickhoff *et al.*, 2005), in SPM12. (L = left hemisphere, R = right hemisphere; x, y, z = co-ordinates of maximum activated voxel in standard MNI152 space, *F* / *t* = *F* / *t* stat at this voxel. (A-G) Peaks are listed at p<0.05 FDR cluster corrected (cluster-forming threshold: p<0.001). *No label in Anatomy toolbox.

| **Cluster** | **Region** | **Hemisphere** | **MNI coordinates** | | |  |
| --- | --- | --- | --- | --- | --- | --- |
|  |  |  | **x** | **y** | **z** | ***F* / *t*** |
| **A** All effects (F test) | | | | | | |
| 1 | Superior Parietal Lobule | R | 15 | -70 | 59 | 36.19 |
|  | Precuneus | L | -12 | -70 | 56 | 34.89 |
|  | Cerebellum (Crus 1) | L | -15 | -88 | -13 | 32.44 |
|  | Lingual Gyrus | L | -15 | -91 | -4 | 31.07 |
|  | Fusiform Gyrus | L | -27 | -49 | -7 | 30.68 |
|  | SupraMarginal Gyrus | R | 51 | -34 | 50 | 28.27 |
|  | Middle Frontal Gyrus | R | 39 | 41 | 26 | 27.51 |
|  | Precentral Gyrus | L | -42 | -13 | 59 | 26.72 |
|  | Lingual Gyrus | R | 18 | -91 | -1 | 26.63 |
|  | Inferior Frontal Gyrus (p. Opercularis) | R | 57 | 11 | 17 | 26.18 |
|  | Occipital Pole* | R | 48 | 14 | -1 | 25.58 |
| 2 | Middle Orbital Gyrus | L | 0 | 56 | -4 | 20.70 |
|  | Anterior Cingulate Cortex | L | 0 | 38 | 2 | 15.43 |
|  | Middle Orbital Gyrus | L | -27 | 35 | -10 | 11.30 |
|  | Superior Medial Frontal Gyrus | L | 0 | 56 | 44 | 10.17 |
|  | Superior Medial Frontal Gyrus | L | 0 | 59 | 38 | 9.69 |
|  | Superior Medial Frontal Gyrus | L | -3 | 62 | 35 | 8.97 |
|  | Frontal Pole* | L | -6 | 68 | 23 | 8.74 |
|  | Frontal Pole* | R | 3 | 71 | 14 | 8.47 |
|  | Subcallosal Cortex* | L | 0 | 11 | -10 | 8.29 |
|  | Middle Orbital Gyrus | R | 0 | 23 | -4 | 7.51 |
|  | Superior Frontal Gyrus | L | -18 | 44 | 50 | 5.07 |
| 3 | Precentral Gyrus | R | 36 | -16 | 68 | 13.12 |
|  | Precentral Gyrus | R | 48 | -16 | 59 | 6.56 |
|  | Postcentral Gyrus | R | 30 | -34 | 74 | 5.50 |
|  | Superior Frontal Gyrus | R | 21 | -16 | 77 | 5.19 |
|  | Precentral Gyrus | R | 51 | -13 | 56 | 4.94 |
|  | Postcentral Gyrus | R | 27 | -40 | 74 | 4.22 |
| 4 | Middle Orbital Gyrus | R | 30 | 35 | -10 | 11.73 |
| **B** Oxytocin Pain Systole | | | | | | |
| 1 | Precuneus | L | -12 | -70 | 56 | 8.49 |
|  | SupraMarginal Gyrus | R | 51 | -34 | 50 | 8.20 |
|  | Lingual Gyrus | R | 18 | -88 | 2 | 7.64 |
|  | Inferior Frontal Gyrus (p. Opercularis) | R | 57 | 11 | 17 | 7.62 |
|  | Frontal Operculum Cortex* | R | 48 | 14 | -1 | 7.58 |
|  | Precuneus | R | 12 | -67 | 59 | 7.55 |
|  | Superior Parietal Lobule | R | 18 | -73 | 56 | 7.49 |
|  | Lingual Gyrus | L | -15 | -91 | -13 | 7.42 |
|  | Middle Occipital Gyrus | L | -15 | -94 | -1 | 7.37 |
|  | Middle Frontal Gyrus | R | 42 | 41 | 23 | 7.28 |
|  | Superior Parietal Lobule | R | 27 | -67 | 56 | 6.97 |
| **C** Oxytocin Pain Diastole | | | | | | |
| 1 | Precuneus | L | -9 | -70 | 59 | 6.13 |
|  | SupraMarginal Gyrus | R | 51 | -34 | 50 | 6.05 |
|  | Precuneus | R | 12 | -70 | 59 | 5.57 |
|  | Postcentral Gyrus | L | -48 | -37 | 56 | 5.45 |
|  | Precentral Gyrus | L | -42 | -22 | 62 | 5.11 |
|  | Inferior Parietal Lobule | L | -39 | -55 | 50 | 4.77 |
|  | Inferior Parietal Lobule | L | -39 | -43 | 41 | 4.11 |
|  | Inferior Parietal Lobule | L | -45 | -40 | 41 | 3.87 |
|  | Angular Gyrus | R | 36 | -61 | 44 | 3.72 |
| 2 | Lingual Gyrus | R | 18 | -88 | 2 | 5.46 |
|  | Cerebellum (VI) | R | 15 | -88 | -10 | 5.01 |
|  | Cerebellum (Crus 1) | R | 36 | -58 | -31 | 4.54 |
|  | Cerebellum (VI) | R | 30 | -61 | -31 | 4.45 |
|  | Cerebellar Vermis (6) | R | 6 | -73 | -16 | 4.06 |
|  | Cerebellar Vermis (7) | R | 3 | -73 | -22 | 3.91 |
|  | Cerebellum* | R | 27 | -49 | -28 | 3.88 |
|  | Cerebellum (IV-V) | R | 9 | -61 | -16 | 3.81 |
|  | Cerebellum* | L | -6 | -67 | -22 | 3.65 |
| 3 | Inferior Frontal Gyrus (p. Orbitalis) | R | 51 | 14 | -1 | 5.44 |
|  | Inferior Frontal Gyrus (p. Opercularis) | R | 57 | 11 | 14 | 5.13 |
|  | Putamen | R | 30 | 2 | 2 | 3.75 |
|  | Insular Cortex* | R | 33 | 11 | 2 | 3.60 |
|  | Pallidum | R | 24 | -10 | 2 | 3.42 |
| 4 | Frontal Operculum Cortex* | L | -48 | 11 | -1 | 4.59 |
|  | Insula Cortex | L | -45 | 2 | 5 | 4.24 |
|  | Insula Cortex | L | -39 | 8 | 2 | 4.14 |
|  | Pallidum* | L | -27 | -7 | -4 | 4.07 |
|  | Putamen* | L | -33 | -1 | -1 | 3.77 |
|  | Inferior Frontal Gyrus (p. Opercularis) | L | -57 | 11 | 14 | 3.68 |
|  | Inferior Frontal Gyrus (p. Opercularis) | L | -60 | 14 | 20 | 3.55 |
| 5 | Frontal Pole* | R | 48 | 44 | -13 | 4.60 |
|  | Middle Frontal Gyrus | R | 45 | 47 | 20 | 4.15 |
|  | Middle Frontal Gyrus | R | 39 | 41 | 26 | 4.03 |
|  | Inferior Frontal Gyrus (p. Triangularis) | R | 51 | 35 | 29 | 3.81 |
|  | Middle Frontal Gyrus | R | 45 | 53 | 8 | 3.73 |
| 6 | Lingual Gyrus | L | -12 | -91 | -13 | 5.03 |
|  | Middle Occipital Gyrus | L | -18 | -94 | 2 | 4.77 |
| 7 | Cerebellum (Crus 1) | L | -42 | -58 | -31 | 5.10 |
|  | Cerebellum (Crus 1) | L | -33 | -58 | -31 | 4.97 |
| 8 | Frontal Pole* | L | -39 | 59 | 2 | 4.19 |
|  | Middle Orbital Gyrus | L | -42 | 53 | -4 | 4.04 |
|  | Middle Orbital Gyrus | L | -45 | 44 | -4 | 4.04 |
| 9 | Middle Frontal Gyrus | L | -33 | 50 | 23 | 4.42 |
|  | Middle Frontal Gyrus | L | -42 | 38 | 32 | 3.37 |
| 10 | Precentral Gyrus | L | -33 | -1 | 65 | 4.06 |
|  | Posterior-Medial Frontal | L | -12 | 5 | 74 | 3.75 |
|  | Superior Frontal Gyrus | L | -18 | 2 | 74 | 3.66 |
| 11 | Posterior-Medial Frontal | L | -6 | -4 | 62 | 4.44 |
|  | Medial Cingulate Cortex | L | -3 | 5 | 47 | 3.24 |
| **D** Oxytocin No Pain Systole | | | | | | |
| 1 | Precuneus | L | -12 | -73 | 56 | 5.48 |
|  | Postcentral Gyrus | L | -45 | -13 | 59 | 5.44 |
|  | Superior Parietal Lobule | L | -24 | -67 | 59 | 4.77 |
|  | Postcentral Gyrus | L | -48 | -37 | 56 | 4.17 |
|  | Inferior Parietal Lobule | L | -48 | -40 | 44 | 4.16 |
|  | Inferior Parietal Lobule | L | -36 | -58 | 56 | 4.15 |
|  | Postcentral Gyrus | L | -48 | -28 | 59 | 4.00 |
|  | Precentral Gyrus | L | -45 | 5 | 53 | 3.58 |
| 2 | Cerebellum (VI) | L | -12 | -85 | -13 | 7.55 |
|  | Lingual Gyrus | L | -12 | -88 | -1 | 7.13 |
|  | Cerebellum (Crus 1) | L | -36 | -55 | -31 | 5.03 |
|  | Cerebellum (Crus 1) | L | -27 | -70 | -25 | 3.60 |
| 3 | Superior Parietal Lobule | R | 15 | -67 | 62 | 6.92 |
|  | Superior Parietal Lobule | R | 15 | -73 | 59 | 6.91 |
|  | SupraMarginal Gyrus | R | 51 | -34 | 47 | 5.59 |
| 4 | Middle Frontal Gyrus | R | 39 | 41 | 26 | 4.98 |
|  | Frontal Pole* | R | 42 | 59 | 2 | 4.14 |
|  | Middle Frontal Gyrus | R | 42 | 29 | 35 | 3.74 |
|  | Middle Orbital Gyrus | R | 45 | 53 | -4 | 3.68 |
|  | Frontal Pole* | R | 45 | 47 | -13 | 3.68 |
|  | Middle Frontal Gyrus | R | 39 | 56 | 17 | 3.41 |
| 5 | Inferior Frontal Gyrus (p. Opercularis) | R | 60 | 11 | 17 | 4.69 |
|  | Insula Cortex | R | 48 | 11 | -1 | 4.65 |
|  | Inferior Frontal Gyrus (p. Opercularis) | R | 57 | 14 | 5 | 3.90 |
|  | Insular Cortex | R | 45 | 2 | 11 | 3.26 |
| 6 | Heschls Gyrus | L | -42 | -25 | 17 | 5.16 |
|  | Middle Temporal Gyrus | L | -57 | -37 | 2 | 4.31 |
|  | Middle Temporal Gyrus* | L | -48 | -34 | -1 | 4.20 |
| 7 | Lingual Gyrus | R | 21 | -88 | 2 | 5.50 |
|  | Cuneus | R | 21 | -94 | 14 | 4.26 |
| 8 | Middle Frontal Gyrus | L | -36 | 50 | 23 | 4.63 |
|  | Frontal Pole* | L | -45 | 50 | -7 | 4.02 |
|  | Frontal Pole* | L | -39 | 59 | 2 | 3.91 |
|  | Frontal Pole* | L | -45 | 50 | 14 | 3.32 |
| 9 | Insular Cortex | L | -45 | -1 | 8 | 4.72 |
|  | Temporal Pole | L | -51 | 14 | -4 | 3.89 |
| 10 | Posterior-Medial Frontal | L | -6 | -4 | 59 | 4.69 |
|  | MCC | L | -6 | 8 | 41 | 3.33 |
| 11 | Insular Cortex* | L | -30 | -4 | -4 | 4.44 |
|  | Putamen | L | -24 | -1 | -1 | 4.24 |
| 12 | Cerebellum (VI) | R | 33 | -52 | -31 | 4.43 |
| **E** Oxytocin No Pain Diastole | | | | | | |
| 1 | Cerebellum (Crus 1) | L | -12 | -85 | -16 | 7.68 |
|  | Lingual Gyrus | L | -9 | -85 | -1 | 6.89 |
|  | Lingual Gyrus | L | -12 | -82 | -4 | 6.88 |
|  | Lingual Gyrus | L | -12 | -91 | -4 | 6.84 |
|  | Superior Occipital Gyrus | L | -18 | -91 | 11 | 6.00 |
|  | Cerebellum (Crus 1) | L | -39 | -58 | -28 | 5.11 |
|  | Cerebellum (Crus 1) | L | -45 | -61 | -28 | 4.97 |
|  | Cerebellum* | L | -18 | -67 | -31 | 4.12 |
| 2 | Precentral Gyrus | L | -42 | -13 | 56 | 5.74 |
|  | Precentral Gyrus | L | -30 | -22 | 68 | 4.89 |
|  | Postcentral Gyrus | L | -48 | -31 | 56 | 4.57 |
|  | Postcentral Gyrus | L | -45 | -40 | 59 | 4.41 |
|  | Precentral Gyrus | L | -30 | -34 | 68 | 4.25 |
|  | Middle Frontal Gyrus | L | -39 | 11 | 53 | 3.85 |
|  | Middle Frontal Gyrus | L | -39 | 14 | 44 | 3.82 |
|  | Middle Frontal Gyrus | L | -45 | 17 | 47 | 3.77 |
|  | Superior Frontal Gyrus | L | -15 | 2 | 74 | 3.74 |
|  | Superior Frontal Gyrus | L | -15 | -7 | 77 | 3.68 |
|  | Middle Frontal Gyrus | L | -36 | 8 | 59 | 3.64 |
| 3 | Precuneus | R | 12 | -70 | 59 | 6.98 |
|  | Superior Parietal Lobule | R | 15 | -76 | 56 | 6.96 |
|  | Precuneus | L | -9 | -70 | 59 | 5.58 |
|  | Precuneus | L | -12 | -73 | 56 | 5.48 |
|  | Superior Parietal Lobule | L | -24 | -67 | 59 | 4.62 |
|  | Inferior Parietal Lobule | L | -39 | -58 | 50 | 3.61 |
|  | Angular Gyrus | L | -36 | -61 | 41 | 3.58 |
|  | Angular Gyrus | R | 42 | -61 | 44 | 3.14 |
| 4 | Middle Frontal Gyrus | R | 39 | 41 | 23 | 5.76 |
|  | Middle Orbital Gyrus | R | 45 | 50 | -7 | 4.90 |
|  | Frontal Pole* | R | 48 | 47 | -10 | 4.88 |
|  | Middle Frontal Gyrus | R | 42 | 56 | 11 | 4.32 |
|  | Middle Frontal Gyrus | R | 33 | 32 | 32 | 4.30 |
|  | Middle Frontal Gyrus | R | 30 | 53 | 29 | 4.29 |
|  | Middle Frontal Gyrus | R | 39 | 59 | 8 | 4.23 |
|  | Middle Orbital Gyrus | R | 45 | 56 | 2 | 4.20 |
| 5 | Middle Frontal Gyrus | L | -30 | 50 | 23 | 4.65 |
|  | Frontal Pole* | L | -42 | 44 | -16 | 4.24 |
|  | Frontal Pole* | L | -45 | 47 | -10 | 4.21 |
|  | Middle Orbital Gyrus | L | -42 | 50 | -7 | 4.21 |
|  | Middle Frontal Gyrus | L | -36 | 56 | 14 | 4.02 |
|  | Frontal Pole* | L | -33 | 62 | -1 | 3.64 |
|  | Middle Frontal Gyrus | L | -33 | 38 | 32 | 3.49 |
| 6 | Temporal Pole | L | -51 | 14 | -4 | 4.85 |
|  | Insular Cortex | L | -45 | -1 | 8 | 4.84 |
|  | Superior Temporal Gyrus | L | -48 | 2 | 2 | 4.70 |
|  | Putamen* | L | -30 | -7 | -4 | 4.01 |
|  | Putamen | L | -24 | -1 | 2 | 3.87 |
|  | Hippocampus | L | -30 | -16 | -10 | 3.44 |
| 7 | Lingual Gyrus | R | 21 | -88 | 2 | 5.73 |
| 8 | Insular Cortex | R | 48 | 8 | -4 | 4.60 |
|  | Inferior Frontal Gyrus (p. Opercularis) | R | 60 | 11 | 14 | 4.51 |
|  | Insular Cortex | R | 48 | 5 | 8 | 4.11 |
| 9 | SupraMarginal Gyrus | R | 51 | -34 | 47 | 4.91 |
|  | SupraMarginal Gyrus | R | 48 | -46 | 50 | 4.54 |
| 10 | Cerebellum (VI) | R | 9 | -64 | -16 | 4.63 |
|  | Cerebellum* | R | 27 | -49 | -28 | 4.49 |
|  | Cerebellum (Crus 1) | R | 39 | -61 | -28 | 4.28 |
|  | Cerebellum* | R | 39 | -55 | -37 | 4.07 |
|  | Cerebellar Vermis (7) | C | 3 | -70 | -22 | 3.90 |
|  | Cerebellum (VI) | R | 30 | -61 | -31 | 3.72 |
|  | Cerebellum* | R | 18 | -55 | -22 | 3.62 |
| 11 | Superior Temporal Gyrus | L | -45 | -28 | 17 | 4.81 |
|  | Insular Cortex* | L | -30 | -25 | 20 | 4.35 |
|  | Heschls Gyrus | L | -33 | -28 | 17 | 4.34 |
| 12 | Posterior-Medial Frontal | L | -6 | -4 | 62 | 4.38 |
|  | MCC | L | -6 | 8 | 41 | 3.37 |
| **F** Placebo Pain Systole | | | | | | |
| 1 | Superior Parietal Lobule | L | -15 | -70 | 56 | 8.37 |
|  | Inferior Frontal Gyrus (p. Opercularis) | R | 54 | 11 | 17 | 8.05 |
|  | Middle Frontal Gyrus | R | 42 | 41 | 23 | 7.35 |
|  | Frontal Operculum Cortex* | R | 48 | 14 | -1 | 7.26 |
|  | SupraMarginal Gyrus | R | 51 | -37 | 50 | 7.24 |
|  | Precentral Gyrus | L | -30 | -4 | 68 | 7.18 |
|  | Insular Cortex | R | 45 | 11 | 2 | 7.16 |
|  | Middle Frontal Gyrus | L | -33 | 53 | 23 | 7.08 |
|  | Middle Frontal Gyrus | L | -27 | -1 | 65 | 7.06 |
|  | Cerebellum (VI) | R | 21 | -85 | -13 | 7.05 |
|  | Inferior Frontal Gyrus (p. Orbitalis) | R | 51 | 17 | -7 | 7.04 |
| **G** Placebo Pain Diastole | | | | | | |
| 1 | Superior Parietal Lobule | L | -15 | -70 | 56 | 7.30 |
|  | Superior Parietal Lobule | R | 15 | -73 | 59 | 6.66 |
|  | Inferior Frontal Gyrus (p. Opercularis) | R | 57 | 11 | 17 | 6.43 |
|  | Middle Frontal Gyrus | L | -30 | -1 | 68 | 6.41 |
|  | Inferior Frontal Gyrus (p. Orbitalis) | R | 51 | 17 | -7 | 6.40 |
|  | Middle Frontal Gyrus | L | -33 | 53 | 23 | 6.25 |
|  | Middle Frontal Gyrus | R | 39 | 41 | 26 | 6.22 |
|  | Middle Frontal Gyrus | R | 30 | 8 | 62 | 6.19 |
|  | Inferior Frontal Gyrus (p. Orbitalis) | L | -48 | 14 | -4 | 6.05 |
|  | Precentral Gyrus | L | -36 | -19 | 62 | 5.87 |
|  | Inferior Parietal Lobule | L | -39 | -49 | 44 | 5.87 |
| 2 | Cerebellum (Crus 1) | R | 33 | -64 | -28 | 6.13 |
|  | Cerebellum (Crus 1) | R | 21 | -85 | -16 | 5.88 |
|  | Cerebellum (Crus 1) | L | -42 | -61 | -28 | 5.59 |
|  | Cerebellum (VI) | R | 12 | -79 | -13 | 5.53 |
|  | Cerebellum* | R | 33 | -52 | -34 | 5.39 |
|  | Inferior Temporal Gyrus | R | 54 | -58 | -19 | 5.30 |
|  | Cerebellum (VIII) | L | -3 | -73 | -28 | 5.20 |
|  | Cerebellum* | R | 36 | -55 | -37 | 5.18 |
|  | Inferior Occipital Gyrus | L | -54 | -67 | -10 | 5.15 |
|  | Middle Occipital Gyrus | L | -15 | -94 | -1 | 5.01 |
|  | Cerebellum (Crus 1) | L | -27 | -70 | -25 | 4.94 |
| **H** Placebo No Pain Systole | | | | | | |
| 1 | Superior Parietal Lobule | R | 15 | -73 | 59 | 7.32 |
|  | Postcentral Gyrus | L | -45 | -13 | 59 | 6.88 |
|  | Precuneus | L | -12 | -73 | 56 | 6.37 |
|  | Middle Frontal Gyrus | R | 30 | 5 | 65 | 5.85 |
|  | Superior Parietal Lobule | L | -24 | -64 | 56 | 5.84 |
|  | Precentral Gyrus | L | -30 | -4 | 68 | 5.49 |
|  | SupraMarginal Gyrus | R | 48 | -37 | 50 | 5.45 |
|  | Postcentral Gyrus | L | -54 | -19 | 50 | 5.35 |
|  | Precentral Gyrus | L | -39 | -4 | 62 | 5.26 |
|  | Inferior Parietal Lobule | L | -42 | -40 | 41 | 5.17 |
|  | Posterior-Medial Frontal | L | 0 | 2 | 59 | 5.13 |
| 2 | Cerebellum (VI) | L | -6 | -82 | -10 | 8.34 |
|  | Lingual Gyrus | L | -12 | -91 | -4 | 7.52 |
|  | Cerebellum (Crus 1) | L | -42 | -61 | -28 | 6.35 |
|  | Lingual Gyrus | R | 18 | -91 | -1 | 5.89 |
|  | Cerebellum (Crus 1) | R | 39 | -64 | -25 | 5.78 |
|  | Superior Occipital Gyrus | R | 24 | -91 | 14 | 5.71 |
|  | Cerebellum (VI) | R | 33 | -52 | -31 | 5.65 |
|  | Cerebellum* | R | 27 | -52 | -28 | 5.50 |
|  | Cerebellum (VIII) | L | -3 | -73 | -28 | 5.41 |
|  | Cerebellum* | R | 36 | -55 | -34 | 5.31 |
| 3 | Middle Frontal Gyrus | R | 39 | 41 | 26 | 5.90 |
|  | Middle Frontal Gyrus | R | 33 | 50 | 26 | 5.14 |
|  | Middle Frontal Gyrus | R | 51 | 17 | 47 | 4.65 |
|  | Middle Frontal Gyrus | R | 42 | 53 | 14 | 4.50 |
|  | Frontal Pole* | R | 48 | 44 | -10 | 4.26 |
|  | Middle Orbital Gyrus | R | 27 | 56 | -7 | 3.60 |
|  | Middle Orbital Gyrus | R | 42 | 56 | -4 | 3.47 |
|  | Frontal Pole* | R | 36 | 59 | -4 | 3.39 |
| 4 | Inferior Frontal Gyrus (p. Orbitalis) | R | 48 | 14 | -4 | 6.01 |
|  | Inferior Frontal Gyrus (p. Orbitalis) | R | 51 | 17 | -7 | 5.93 |
|  | Inferior Frontal Gyrus (p. Opercularis) | R | 57 | 11 | 17 | 5.26 |
|  | Inferior Frontal Gyrus (p. Orbitalis) | R | 39 | 23 | -13 | 4.68 |
|  | Insular Cortex | R | 33 | 20 | 8 | 4.37 |
| 5 | Inferior Frontal Gyrus (p. Orbitalis) | L | -45 | 14 | -4 | 5.16 |
|  | Inferior Frontal Gyrus (p. Opercularis) | L | -60 | 8 | 20 | 3.92 |
|  | Insular Cortex | L | -42 | 2 | 8 | 3.88 |
|  | Putamen* | L | -33 | -1 | 5 | 3.82 |
|  | Inferior Frontal Gyrus (p. Triangularis) | L | -57 | 14 | 17 | 3.82 |
|  | Inferior Frontal Gyrus (p. Triangularis) | L | -57 | 23 | 23 | 3.57 |
|  | Temporal Pole | L | -54 | 11 | -13 | 3.48 |
| 6 | Middle Frontal Gyrus | L | -33 | 50 | 20 | 5.15 |
| 7 | Superior Temporal Gyrus | L | -45 | -31 | 17 | 4.40 |
|  | Planum Temporale* | L | -39 | -31 | 8 | 3.25 |
| **I** Placebo No Pain Diastole | | | | | | |
| 1 | Superior Parietal Lobule | R | 15 | -70 | 62 | 7.67 |
|  | Postcentral Gyrus | L | -45 | -13 | 56 | 7.47 |
|  | Middle Frontal Gyrus | R | 42 | 41 | 26 | 6.99 |
|  | Middle Frontal Gyrus | R | 33 | 53 | 29 | 6.55 |
|  | Superior Frontal Gyrus | R | 27 | 8 | 65 | 6.43 |
|  | Superior Parietal Lobule | L | -15 | -70 | 56 | 6.34 |
|  | Precentral Gyrus | L | -33 | -4 | 65 | 6.25 |
|  | Middle Frontal Gyrus | L | -33 | 50 | 23 | 6.24 |
|  | Superior Parietal Lobule | L | -24 | -67 | 59 | 6.10 |
|  | Precentral Gyrus | L | -39 | -4 | 62 | 6.04 |
|  | SupraMarginal Gyrus | R | 48 | -49 | 47 | 6.02 |
| 2 | Cerebellum (VI) | L | -9 | -85 | -10 | 8.19 |
|  | Lingual Gyrus | L | -12 | -82 | -1 | 8.13 |
|  | Lingual Gyrus | R | 18 | -91 | -1 | 6.28 |
|  | Middle Occipital Gyrus | L | -21 | -91 | 14 | 6.17 |
|  | Superior Occipital Gyrus | R | 24 | -91 | 14 | 6.09 |
|  | Cerebellum (VI) | R | 18 | -88 | -10 | 5.95 |
|  | Cerebellum (Crus 1) | L | -42 | -64 | -22 | 5.68 |
|  | Cerebellum (Crus 1) | L | -42 | -58 | -28 | 5.68 |
|  | Inferior Temporal Gyrus | R | 60 | -52 | -13 | 5.46 |
|  | Cerebellum (Crus 1) | L | -27 | -70 | -25 | 5.42 |
|  | Cerebellum (VI) | R | 27 | -58 | -28 | 5.36 |
| 3 | Inferior Frontal Gyrus (p. Orbitalis) | R | 48 | 14 | -4 | 5.63 |
|  | Inferior Frontal Gyrus (p. Opercularis) | R | 60 | 11 | 11 | 4.76 |
|  | Inferior Frontal Gyrus (p. Opercularis) | R | 57 | 11 | 17 | 4.65 |
|  | Inferior Frontal Gyrus (p. Orbitalis) | R | 39 | 23 | -13 | 4.29 |
| 4 | Rolandic Operculum | L | -42 | -28 | 17 | 4.88 |
|  | Middle Temporal Gyrus | L | -66 | -31 | 5 | 4.12 |
|  | Middle Temporal Gyrus* | L | -45 | -34 | -4 | 4.11 |
|  | Middle Temporal Gyrus | L | -60 | -37 | 5 | 4.05 |
|  | Middle Temporal Gyrus | L | -54 | -40 | 2 | 4.02 |
|  | Superior Temporal Gyrus | L | -48 | -37 | 11 | 3.89 |
|  | Middle Temporal Gyrus* | L | -48 | -46 | -1 | 3.52 |
| 5 | Putamen* | L | -24 | 14 | 8 | 4.58 |
|  | Caudate* | L | -21 | 14 | -7 | 3.80 |
|  | Pallidum | L | -18 | 2 | 2 | 3.74 |
|  | Putamen | L | -30 | -10 | 2 | 3.49 |
|  | Putamen* | L | -27 | 8 | -4 | 3.44 |
| 6 | Inferior Frontal Gyrus* | L | -54 | 14 | -1 | 4.40 |
|  | Inferior Frontal Gyrus (p. Orbitalis) | L | -51 | 17 | -4 | 4.32 |
|  | Frontal Operculum Cortex* | L | -48 | 11 | -1 | 4.20 |
|  | Inferior Frontal Gyrus* | L | -57 | 14 | 5 | 4.17 |
|  | Temporal Pole | L | -48 | 5 | -1 | 4.16 |
|  | Insular Cortex | L | -42 | -4 | 11 | 4.03 |
|  | Inferior Frontal Gyrus* | L | -60 | 14 | 17 | 3.95 |
|  | Inferior Frontal Gyrus (p. Opercularis) | L | -60 | 2 | 17 | 3.93 |
| 7 | Thalamus | L | -18 | -22 | 11 | 5.09 |
| 8 | Caudate Nucleus | R | 18 | 20 | 8 | 4.68 |
|  | Insular Cortex* | R | 30 | 20 | 8 | 3.54 |
| 9 | Frontal Orbital Cortex* | R | 21 | 20 | -7 | 3.28 |
| **J** Placebo > Oxytocin | | | | | | |
| 1 | Middle Temporal Gyrus | R | 48 | -64 | 5 | 5.64 |
|  | Fusiform Gyrus | R | 42 | -76 | -13 | 5.60 |
|  | Inferior Temporal Gyrus* | R | 48 | -46 | -4 | 5.48 |
|  | Lateral Occipital Cortex* | R | 36 | -64 | 2 | 5.16 |
|  | Lateral Occipital Cortex* | R | 30 | -64 | 29 | 4.61 |
|  | Inferior Temporal Gyrus | R | 54 | -34 | -10 | 4.56 |
|  | Precuneus | R | 21 | -64 | 32 | 4.53 |
|  | Middle Temporal Gyrus | R | 42 | -67 | 14 | 4.36 |
|  | Calcarine Gyrus | R | 24 | -64 | 14 | 4.36 |
|  | Cuneus | R | 18 | -82 | 41 | 4.19 |
|  | Cerebellum (VII) | R | 33 | -73 | -40 | 4.05 |
| 2 | Lateral Occipital Cortex* | L | -39 | -61 | 2 | 6.60 |
|  | Inferior Temporal Gyrus | L | -45 | -70 | -1 | 6.21 |
|  | Fusiform Gyrus | L | -36 | -49 | -7 | 5.40 |
|  | Middle Occipital Gyrus | L | -36 | -79 | 26 | 4.79 |
|  | Cerebral White Matter* | L | -30 | -52 | 20 | 4.69 |
|  | Cerebellum (IV-V) | L | -18 | -46 | -13 | 4.65 |
|  | Middle Occipital Gyrus | L | -45 | -85 | 8 | 4.58 |
|  | Middle Occipital Gyrus | L | -39 | -82 | 14 | 4.37 |
|  | Precuneus Cortex* | L | -27 | -58 | 14 | 4.26 |
| 3 | Posterior-Medial Frontal | R | 12 | 5 | 71 | 5.23 |
|  | Middle Frontal Gyrus | R | 30 | 8 | 62 | 4.10 |
|  | Middle Frontal Gyrus* | R | 30 | 5 | 47 | 4.09 |
|  | Middle Frontal Gyrus | R | 30 | 8 | 53 | 4.02 |
|  | Superior Frontal Gyrus | R | 24 | -1 | 59 | 4.02 |
|  | Precentral Gyrus | R | 33 | -4 | 53 | 3.57 |
| 4 | Posterior-Medial Frontal | L | -3 | 17 | 50 | 4.38 |
|  | Posterior-Medial Frontal | L | -6 | 14 | 53 | 4.34 |
|  | Superior Medial Gyrus | L | -3 | 23 | 47 | 4.26 |
|  | Superior Medial Gyrus | R | 9 | 29 | 56 | 4.12 |
|  | MCC | R | 15 | 17 | 38 | 4.08 |
|  | Posterior-Medial Frontal | R | 9 | 17 | 50 | 3.90 |
|  | MCC | L | -12 | 23 | 38 | 3.66 |
|  | Superior Medial Gyrus | L | -9 | 29 | 41 | 3.55 |
|  | Superior Frontal Gyrus | R | 18 | 23 | 56 | 3.50 |
|  | Superior Medial Gyrus | R | 9 | 23 | 65 | 3.45 |
| 5 | Paracingulate Gyrus* | L | -15 | 38 | 14 | 4.96 |
|  | ACC | L | -9 | 38 | 17 | 4.84 |

Barr, D.J., Levy, R., Scheepers, C. & Tily, H.J. (2013) Random effects structure for confirmatory hypothesis testing: Keep it maximal. *J Mem Lang,* 68 (3).

de Haan, H.A., van der Palen, J., Wijdeveld, T.G., Buitelaar, J.K. & De Jong, C.A. (2014) Alexithymia in patients with substance use disorders: state or trait? *Psychiatry Res,* 216 (1)**,** 137-45.

Eickhoff, S.B., Stephan, K.E., Mohlberg, H., Grefkes, C., Fink, G.R., Amunts, K. & Zilles, K. (2005) A new SPM toolbox for combining probabilistic cytoarchitectonic maps and functional imaging data. *Neuroimage,* 25 (4)**,** 1325-35.

Helling, J. (2009) Non-declarative representational and regulatory systems in alexithymia. *J Trauma Dissociation,* 10 (4)**,** 469-87.

Kagerbauer, S.M., Martin, J., Schuster, T., Blobner, M., Kochs, E.F. & Landgraf, R. (2013) Plasma oxytocin and vasopressin do not predict neuropeptide concentrations in human cerebrospinal fluid. *J Neuroendocrinol,* 25 (7)**,** 668-73.

Landgraf, R. (1985) Plasma oxytocin concentrations in man after different routes of administration of synthetic oxytocin. *Exp Clin Endocrinol,* 85 (2)**,** 245-8.

Little, R.J. (1988) Missing-data adjustments in large surveys. *Journal of Business & Economic Statistics,* 6 (3)**,** 287-296.

Luminet, O., Grynberg, D., Ruzette, N. & Mikolajczak, M. (2011) Personality-dependent effects of oxytocin: greater social benefits for high alexithymia scorers. *Biol Psychol,* 87 (3)**,** 401-6.

Striepens, N., Kendrick, K.M., Hanking, V., Landgraf, R., Wullner, U., Maier, W. & Hurlemann, R. (2013) Elevated cerebrospinal fluid and blood concentrations of oxytocin following its intranasal administration in humans. *Sci. Rep.,* 3.

Van Buuren, S. & Groothuis-Oudshoorn, K., (2011) Mice:Multivariate Imputation by Chained Equations in R. Journal of Statistical Software, 1-67.

Wotjak, C.T., Ganster, J., Kohl, G., Holsboer, F., Landgraf, R. & Engelmann, M. (1998) Dissociated central and peripheral release of vasopressin, but not oxytocin, in response to repeated swim stress: new insights into the secretory capacities of peptidergic neurons. *Neuroscience,* 85 (4)**,** 1209-22.
